# Systemic gene therapy with thymosin β4 alleviates glomerular injury in mice

**DOI:** 10.1101/2021.05.24.445384

**Authors:** William J Mason, Daniyal J Jafree, Gideon Pomeranz, Maria Kolatsi-Joannou, Sabrina Pacheco, Dale A Moulding, Anja Wolf, Christian Kupatt, Claire Peppiatt-Wildman, Eugenia Papakrivopoulou, Paul R Riley, David A Long, Elisavet Vasilopoulou

**Author notes:** **Corresponding Author**: Dr Elisavet Vasilopoulou, Medway School of Pharmacy, Division of Natural Sciences, University of Kent, Chatham, Kent, UK.

## Abstract

Plasma ultrafiltration in the kidney occurs across glomerular capillaries, which are surrounded by epithelial cells called podocytes. Podocytes have a unique shape maintained by a complex cytoskeleton, which becomes disrupted in glomerular disease resulting in defective filtration and albuminuria. Lack of endogenous thymosin β4 (TB4), an actin sequestering peptide, exacerbates glomerular injury and disrupts the organisation of the podocyte actin cytoskeleton, however, the effect of exogenous TB4 therapy on podocytopathy is unknown. Here, through interrogating single-cell RNA-sequencing data of isolated glomeruli we demonstrate that Adriamycin, a toxin which injures podocytes and leads to leakage of albumin in the urine of mice, results in reduced levels of podocyte TB4. Systemic administration of an adeno-associated virus vector encoding TB4 prevented Adriamycin-induced podocyte loss and albuminuria. Adriamycin injury was associated with disorganisation of the actin cytoskeleton *in vitro*, which was ameliorated by exogenous TB4. Furthermore, Adriamycin administration in mice was associated with increased prevalence of podocyte vesicles, a mechanism by which albumin may leak into the urine, which was also prevented by TB4. Collectively, we propose that TB4 gene therapy prevents podocyte injury and maintains glomerular filtration via modulation of the podocyte cytoskeleton thus presenting a novel treatment strategy for glomerular disease.

## Introduction

One in ten people worldwide have chronic kidney disease (CKD).^1^ A subset of patients with CKD progress to end-stage kidney disease (ESKD), which requires dialysis or transplantation and is a risk factor for cardiovascular disease and all-cause mortality.^1^ CKD progression is linked to breakdown of the glomerular filtration barrier, the site of ultrafiltration in the kidney, which consists of endothelial cells, the glomerular basement membrane (GBM) and epithelial podocytes.^2,3^ Podocytes have a unique architecture with foot processes that extend from their cell bodies, interdigitate and form slit diaphragms facilitating size and charge-selective filtration and preventing the loss of plasma proteins.^4,5^ In health, podocyte shape is maintained by a complex, highly regulated actin cytoskeleton, which supports the foot processes,^6,7^ and anchors the cell to the GBM.^8^ During glomerular disease, the podocyte cytoskeleton becomes disorganised often leading to podocyte loss accompanied by impaired filtration and leakage of plasma proteins, such as albumin, into the urine.^6,9–11^ Albuminuria is a hallmark of glomerular disease, irrespective of the underlying aetiology.^12^ Therefore, therapies that protect the podocyte cytoskeleton could provide a novel strategy to preserve the integrity of the glomerular filtration barrier, prevent albuminuria and improve glomerular disease progression.

Thymosin β4 (TB4) sequesters monomeric G-actin in mammalian cells^13,14^ and maintains high concentrations of G-actin available for polymerisation into actin filaments (F-actin).^15^ Our previous work has shown that endogenous TB4 is expressed in podocytes and plays a protective role in glomerular disease. We found that lack of endogenous TB4 worsens albuminuria, renal function and glomerular injury in a mouse model of glomerulonephritis, concomitant with redistribution of podocytes from the glomerular tuft, where they contribute to filtration barrier integrity, to the Bowman’s capsule. Furthermore, a direct role for endogenous TB4 on the podocyte cytoskeleton was established *in vitro*, with a shift from cortical actin to cytoplasmic actin stress fibres and enhanced migration observed in podocytes lacking TB4.^16^

These findings raise the possibility that exogenous TB4 could be used as a therapy to protect the podocyte cytoskeleton and slow the progression of glomerular disease. Exogenous TB4 has already shown promise as a treatment for a diverse range of conditions, such as myocardial infarction,^17^ dry eye syndrome,^18^ stroke^19^ and inflammatory lung disease.^20^ Furthermore, exogenous TB4 is protective in several animal models of kidney injury,^21^ including diabetic nephropathy,^22^ unilateral ureteral obstruction^23,24^ and acute ischaemia reperfusion injury,^25^ but none of these studies examined the specific effect of exogenous TB4 on podocytes. Additionally, the studies above administered TB4 peptide, which has a relatively short half-life with enhanced levels in the plasma evident for only 6 hours following injection.^26^ To overcome the rapid metabolic turnover of TB4, recombinant adeno-associated virus (AAV)-mediated gene therapy provides a solution through its ability to induce stable, long term transgene expression via a single injection.^27–29^ Indeed, TB4-encoding AAV constructs have been successfully used for tissue-specific (muscle, heart) or systemic upregulation of TB4 in mouse, rabbit and pig disease models with therapeutic effects.^30–32^

In this study, we show that podocyte injury induced by a toxin (Adriamycin (ADR)) is associated with reduced expression of endogenous TB4 through interrogating single-cell RNA sequencing (scRNAseq) data of isolated glomeruli. Using systemic gene therapy, upregulation of the plasma concentration of TB4 was able to prevent ADR-induced albuminuria and podocyte loss. Further examination of the podocyte F-actin structures *in vitro* revealed that exogenous TB4 prevented ADR-induced cytoskeletal disorganisation. Collectively, our work shows that gene therapy-mediated systemic administration of TB4 presents a novel treatment strategy to protect podocytes from injury and preserve the integrity of the glomerular filtration barrier.

## Results

### ADR is associated with the downregulation of TB4 in podocytes

To induce glomerular damage *in vivo*, we administered ADR, a chemotherapeutic drug, to male BALB/c mice. In BALB/c mice ADR replicates some of the features of human focal segmental glomerulosclerosis (FSGS)^33^ and has toxic effects on podocytes *in vitro* and *in vivo*, including cytoskeletal disorganisation, foot process effacement and a loss of podocyte viability^9,34,35^. Firstly, we examined if ADR administration altered glomerular TB4 expression. To do this, we isolated glomeruli from ADR-administered BALB/c mice by Dynabead perfusion^36^ and using quantitative real-time PCR (qRT-PCR), we found that there was a downregulation of the mRNA transcript for TB4, *Tmsb4x*, at 7 days post ADR administration compared with saline injected mice (P=0.076) but not at 14 days (**Figure 1a**). To further interrogate the expression of *Tmsb4x* in specific glomerular cells following ADR injury, we analysed a published scRNAseq dataset obtained from glomeruli isolated by Dynabeads of healthy or ADR-injured C57BL/6J mice^37^. Using unsupervised clustering analysis we derived ten transcriptionally distinct cell types (**Figure 1b**), which using established markers of glomerular cell types, we identified to represent all the known glomerular cell types (**Supplementary Figure 1a**). Confirming the identity of podocytes, the podocyte cluster expressed both nephrin (Nphs1) and podocin (Nphs2), components of the podocyte slit diaphragm (**Figure 1c**). Comparison of control and ADR-injured glomeruli showed representation of all cell types in both groups (**Supplementary Figure 1b**). As previously described^37^, we found a reduction in the proportion of podocyte cells in the ADR-injured glomeruli compared with controls (**Supplementary Figure 1c**). ADR injury was associated with significant downregulation of *Tmsb4x* in the glomerular tuft, assessed by grouped analysis of glomerular endothelial, mesangial and podocyte cells (**Figure 1d**; 0.14 log fold change; P<0.05). When podocyte cells were analysed individually, we found a larger 0.37 log fold reduction in *Tmsb4x* in those obtained from ADR-injured compared with healthy glomeruli (**Figure 1e**; P<0.05). Collectively, these findings demonstrate that ADR injury is associated with reduced *Tmsb4x* expression in the glomerulus and specifically in podocytes.

**Figure 1.**
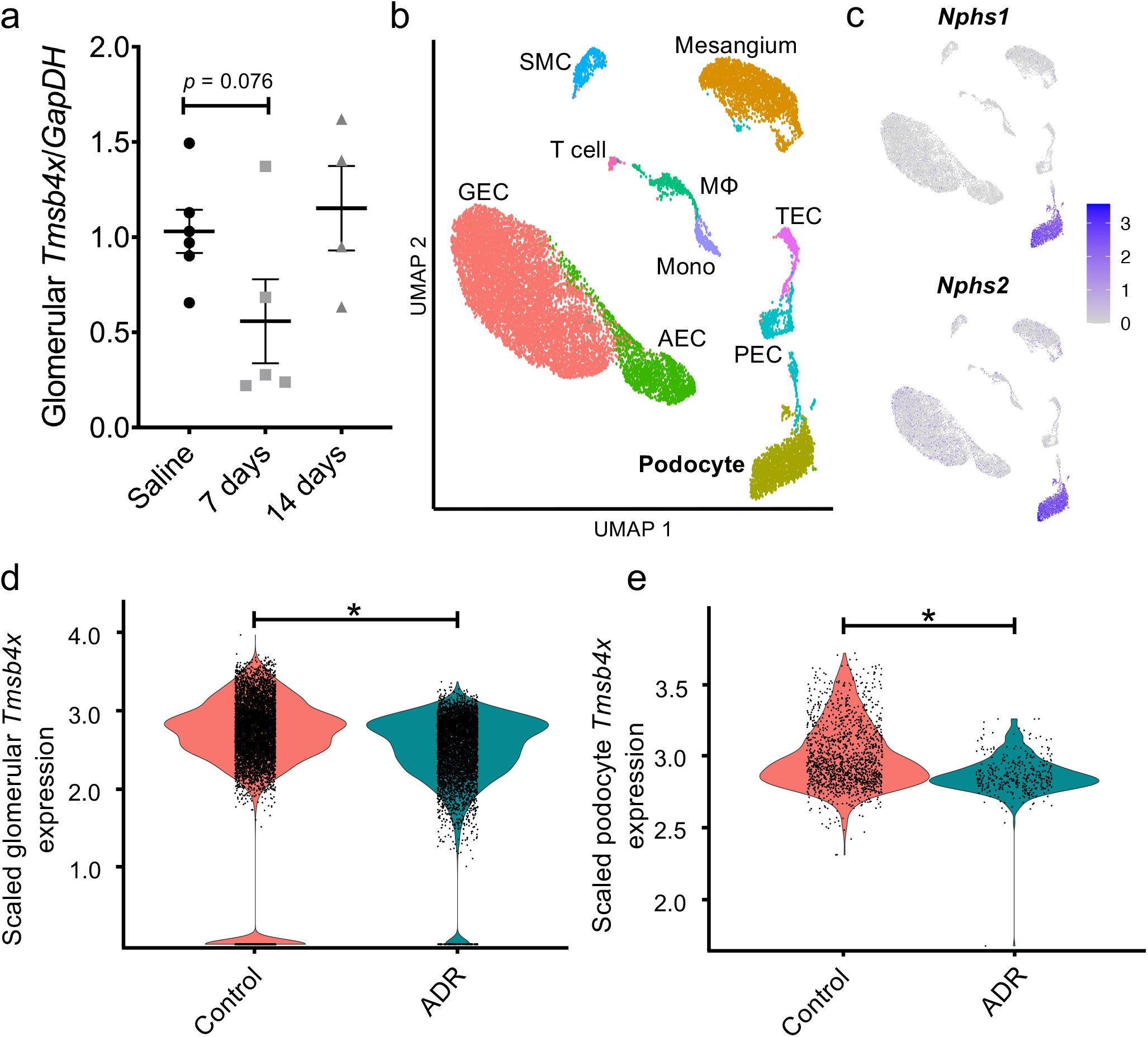
Podocyte-specific *Tmsb4x* is downregulated in murine Adriamycin nephropathy. **(a)** Expression of *Tmsb4x* as assessed by quantitative reverse transcriptase polymerase chain reaction in glomeruli of mice administered with 10 mg/kg of Adriamycin (ADR). **(b)** Uniform manifold approximation and projection (UMAP) from single-cell RNA sequencing data of 8,412 glomerular cells from two wildtype (control) mice and 8,296 glomerular cells from mice with Adriamycin nephropathy (ADR). After analysis and cell type assignment, ten transcriptionally distinct cell populations were discriminated including arterial endothelial cells (AEC), glomerular endothelial cells (GEC), macrophages (MΦ), mesangial cells, monocytes (Mono), parietal epithelial cells (PEC), podocytes, smooth muscle cells (SMC), T cells, tubular epithelial cells (TEC). The markers used for cell type identification and assignment, and the numbers of cell types per condition, are shown in **Figure S1. (c)** Feature plots showing expression of nephrin (Nphs1) and podocin (Nphs2), canonical markers of podocytes, across the dataset. **(d)** Violin plot comparing the scaled expression of *Tmsb4x* of all glomerular cells (podocytes, GECs, mesangium) between ADR (n = 5,602 cells) and control (n = 7,190 cells). An average log fold decrease of 0.13 was detected in ADR compared to control (*: adjusted p value < 0.0001). **(e)** Violin plot comparing the scaled expression of *Tmsb4x* of podocytes between ADR (n = 378 cells) and control (n = 1,486 cells). An average log fold decrease of 0.36 was detected in ADR compared to control (*: adjusted p value < 0.0001).

### Systemic upregulation of TB4 using gene therapy

We then assessed the effects of exogenous TB4 administration on ADR-induced glomerular disease. We took a preventative strategy and BALB/c mice were administered intravenously AAV serotype 2/7 expressing either *Tmsb4x* or the *LacZ* gene as a control.^30,32^ Three weeks after AAV injection ADR was administered (day 0). Fourteen days later the animals were euthanised and we compared mice administered either: (i) AAV-*Tmsb4x* and ADR (*Tmsb4x*/ADR); (ii) AAV-*LacZ* and ADR (*LacZ*/ADR) and (iii) AAV-*LacZ* and saline as our control group (*LacZ*/saline) (**Figure 2a**). Initially, we examined if AAV-*Tmsb4x* altered *Tmsb4x* mRNA levels in the liver, as this is the primary site of transduction following systemic injection of AAV 2/7.^38^ We found a 10-fold increase in liver *Tmsb4x* mRNA levels in the *Tmsb4x*/ADR group compared with the *LacZ*/ADR group (P<0.001) (**Figure 2b**). This corresponded with strong positive immunostaining for TB4 in the liver in *Tmsb4x*/ADR mice (**Figure 2c**). Additionally, as TB4 is a secreted peptide^39^, we measured circulating levels using ELISA and found a 2-fold increase in the *Tmsb4x*/ADR group compared with either the *LacZ*/ADR (P<0.05) or *LacZ*/saline group (P<0.01) (**Figure 2d**). Using qRT-PCR on whole kidney lysates we found no difference in *Tmsb4x* mRNA levels between the three experimental groups (**Figure 2e**) in agreement with previous reports that AAV 2/7 does not specifically target the kidney.^38^

**Figure 2.**
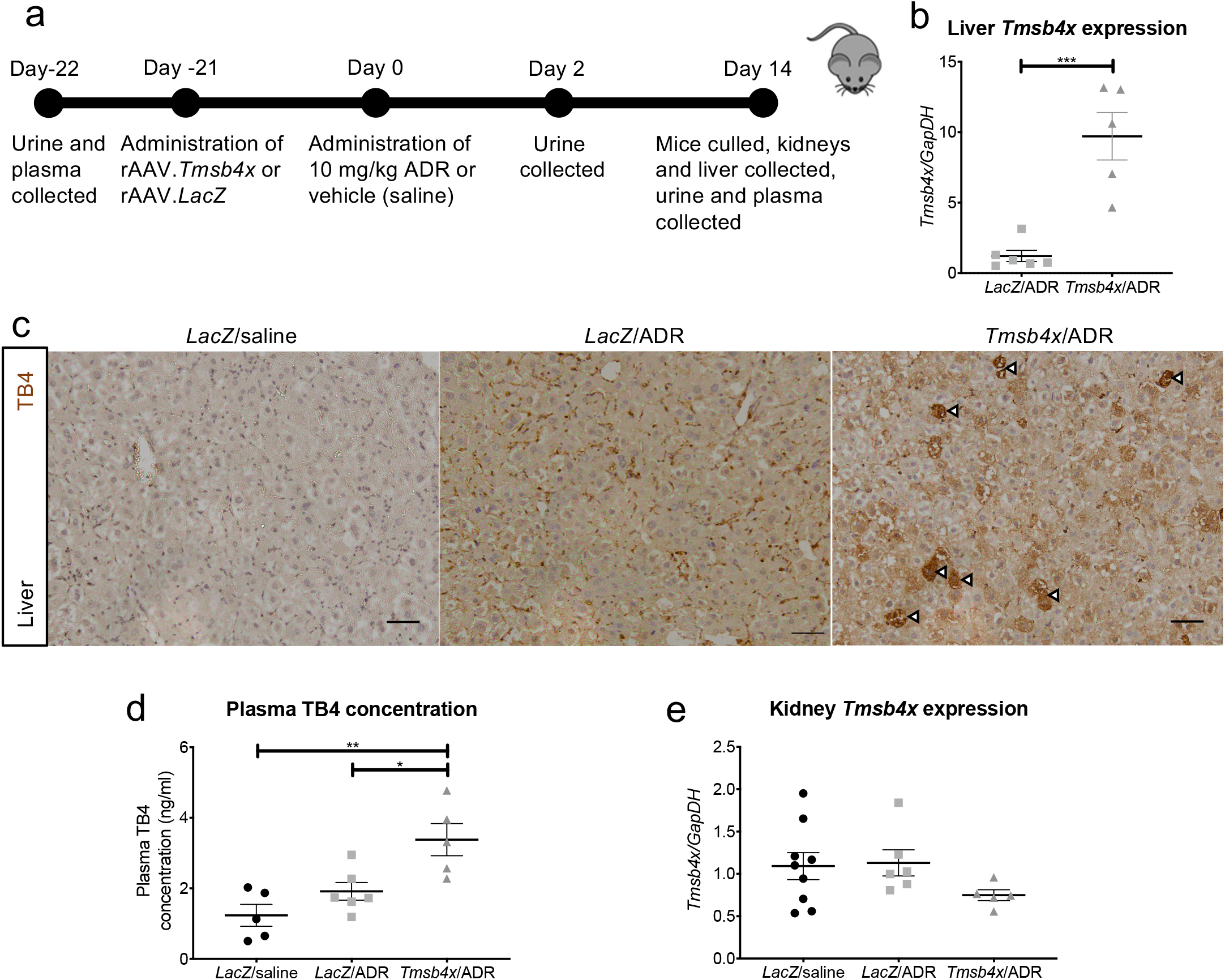
AAV2/7 causes systemic upregulation of thymosin β4 *in vivo*. **(a)** BALB/c mice were injected with AAV-*LacZ* or AAV-*Tmsb4x* three weeks prior to 10 mg/kg ADR or vehicle (saline) administration and culled 14 days after ADR/saline injection. **(b)** Liver expression of *Tmsb4x* mRNA. **(c)** Expression of TB4 peptide in the liver. Arrowheads refer to cells with particularly high expression of TB4. Scale bars = 50 µm. **(d)** Plasma concentration of TB4. **(e)** Whole kidney expression of *Tmsb4x* gene. All experiments were repeated a minimum of 5 times. For gene analysis and plasma concentration, assays were performed in duplicate and the data are presented as the mean±SEM. \**P* ≤ 0.05, \*\**P* ≤ 0.01, ***P≤ 0.001. TB4, *Tmsb4x*, thymosin β4; ADR, Adriamycin; *LacZ*, β-Galactosidase; AAV, adeno-associated virus.

### TB4 prevents ADR-induced albuminuria

Next, we examined the effect of TB4 administration on ADR injury *in vivo*. Firstly, as an overall measure of health, we weighed the mice throughout the time course of the experiment. All mice administered ADR lost a significantly greater proportion of their body weight 48 hours following ADR injection compared with *LacZ*/saline animals (**Figure 3a**). This difference was sustained at both day 7 and 14 after ADR injection. At all time-points there was no difference between ADR mice administered AAV-*LacZ* and AAV-*Tmsb4x*.

**Figure 3.**
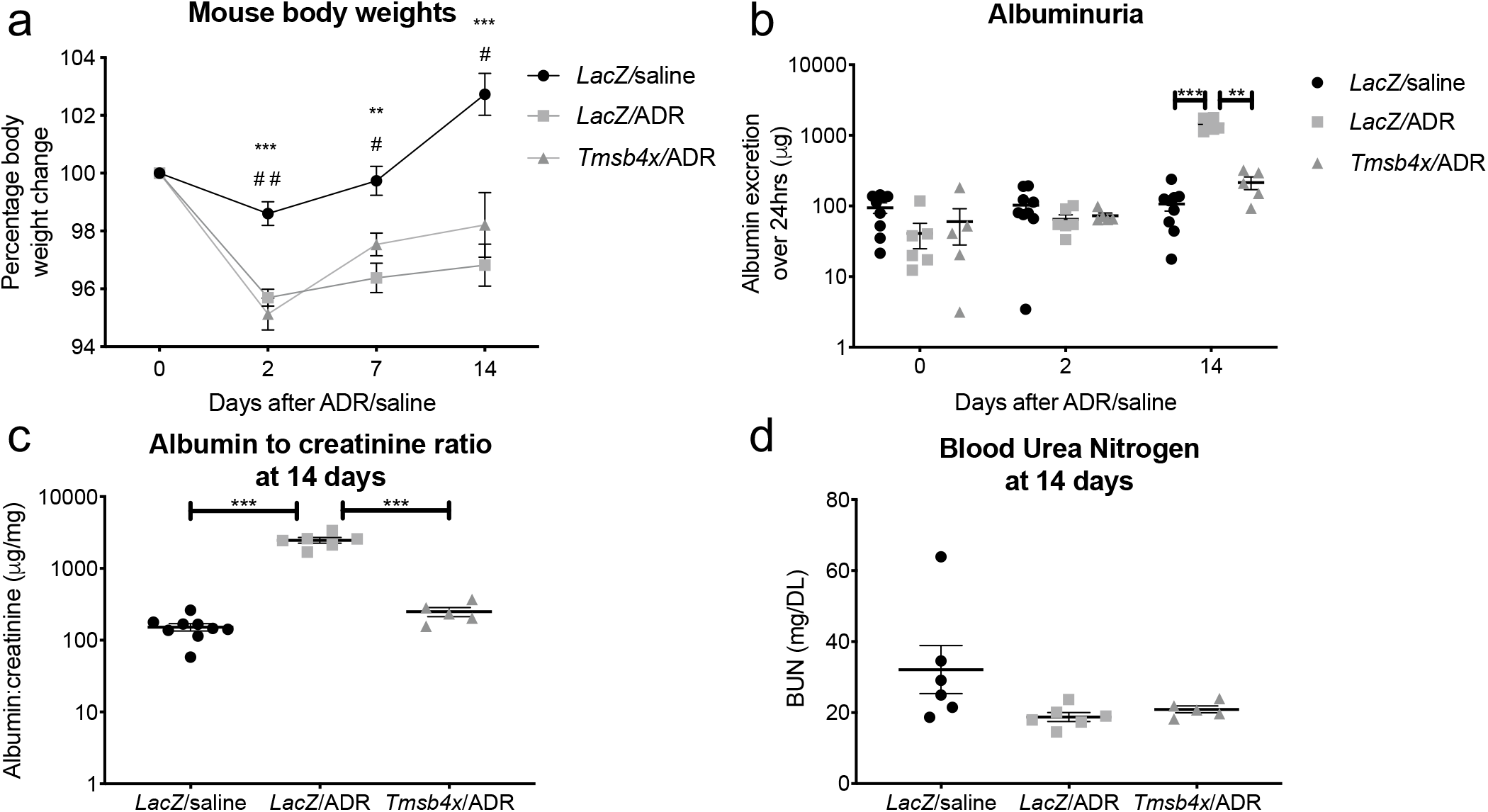
Renal function following ADR/TB4 treatment. **(a)** Mouse weight change 2, 7 and 14 days after ADR/vehicle injection. Statistical annotations using * refer to comparison between *LacZ*/saline & *LacZ*/ADR. Annotations using # refer to comparison between *LacZ*/saline & *Tmsb4x*/ADR. **(b)** Twenty-four hour urinary albumin excretion 0, 2 and 14 days after ADR/vehicle injection. **(c)** Urinary albumin to creatinine ratio 14 days after ADR/vehicle injection. **(d)** Blood urea nitrogen concentration 14 days after ADR/vehicle injection. \**P* ≤ 0.05, \*\**P* ≤ 0.01, ***P≤ 0.001. TB4, *Tmsb4x*, thymosin β4; ADR, Adriamycin; *LacZ*, β-Galactosidase.

We then analysed the levels of albumin in the urine, a marker of glomerular filtration barrier integrity.^40^ Prior to ADR injection, there was no difference in the levels of urinary albumin (*LacZ*/saline group: 95 ±16 μg/24 hours, *LacZ*/ADR group: 41 ±16 μg/24 hours, *Tmsb4x*/ADR group: 60 ±32 μg/24 hours). Similar levels were found two days after ADR administration with no differences between the three groups of mice (*LacZ*/saline group: 103 ±20 μg/24 hours, *LacZ*/ADR group: 65 ±11 μg/24 hours, *Tmsb4x*/ADR group: 73 ±7 μg/24 hours). Conversely, at 14 days post ADR injection, albuminuria significantly increased to 1448 ±115 μg/24 hours in the *LacZ*/ADR mice compared with 107 ±22 μg/24 hours in *LacZ*/saline animals (P<0.001). Strikingly, gene therapy with TB4 prevented this increase with a urinary albumin excretion of 214 ±43 μg/24 hours in the *Tmsb4x*/ADR group (P<0.01 versus *LacZ*/ADR group) (**Figure 3b**). At 14 days, the albumin to creatinine ratio of the *LacZ*/ADR group was also increased 25-fold in comparison with *LacZ*/saline mice (P<0.001). This increase was prevented in the ADR mice treated with TB4 (P<0.001 versus *LacZ*/ADR group; **Figure 3c**). Urea is freely filtered through the glomerular filtration barrier and increased plasma blood urea nitrogen (BUN) concentration is an indicator of declining kidney function^41^. However, we found that there were no significant changes between any of the groups 14 days after ADR administration (**Figure 3d**).

Previously the protective effect of TB4 in CKD has been linked to its anti-inflammatory properties ^16,24,42^. Therefore, we quantified F4/80^+^ macrophage numbers within and around the glomerular tuft by immunohistochemistry (**Supplementary Figure 2a**). F4/80^+^ macrophages within the glomerular tuft were rare across all three groups (**Supplementary Figure 2b**). The number of macrophages outside the glomerular tuft was similar among the groups (**Supplementary Figure 2c**), excluding a role for inflammation in our model and suggesting that the protective effect of TB4 in ADR-induced glomerular injury occurs by other means.

### TB4 prevents ADR-induced podocyte injury *in vivo*

Next, we assessed the effect of exogenous TB4 on podocytes in this model. As ADR has been shown to cause podocyte loss^43^ we quantified the number of Wilms Tumour 1 positive (WT1^+^) podocytes in the glomerulus^44^ (**Figure 4a**). When analysing the results obtained per mouse, we found lower podocyte number and density (WT1^+^ podocytes/glomerular area) in the *LacZ*/ADR group compared with the *LacZ*/saline and *Tmsb4x*/ADR groups, however this was not statistically significant (**Figure 4b, c**). Since one of the features of FSGS is that not all glomeruli are affected to the same extent^45^, we also examined podocyte number at the level of individual glomeruli by assessing a total of at least 250 glomeruli from 5 different mice in each group. Here, we found a significant decrease in podocyte number (8.85 ±0.31 podocytes) and density (5.31×10^−3^ ±0.18⨯10^−3^ podocytes/μm^2^) in the *LacZ*/ADR group compared with *LacZ*/saline (10.94 ±0.25 podocytes and 6.59⨯10^−3^ ±0.13⨯10^−3^ podocytes/μm^2^ respectively) (P<0.001 in both cases). TB4 treatment prevented the ADR-induced podocyte loss and preserved podocyte number (10.38 ±0.34 podocytes) (P<0.001 versus *LacZ*/ADR group) and podocyte density (6.57⨯10^−3^ ±0.20 podocytes/μm^2^) (P<0.001 versus *LacZ*/ADR group) (**Figure 4d, e**).

**Figure 4.**
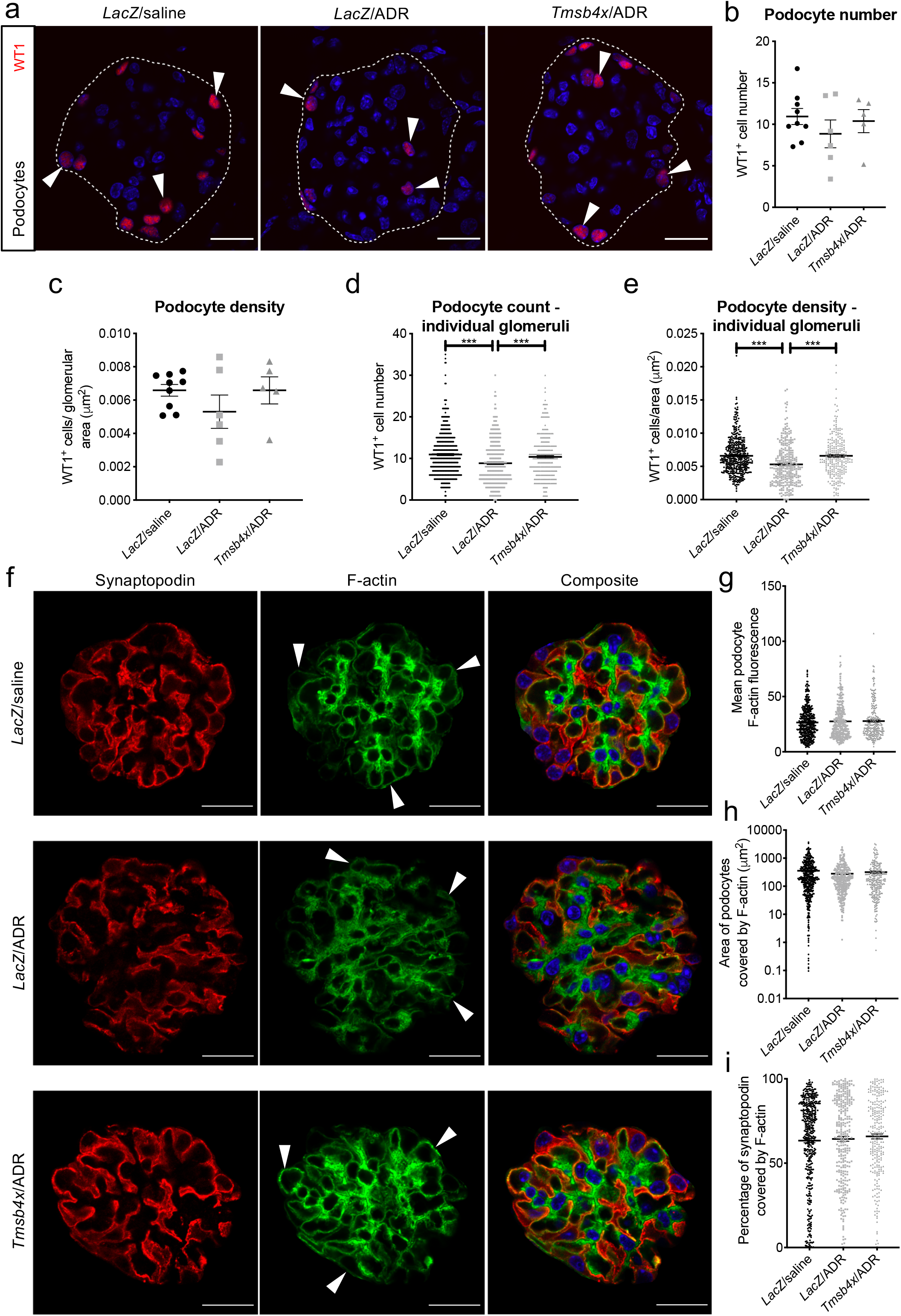
Analysis of podocytes *in vivo* after ADR/TB4 treatment. **(a)** Representative images of WT1^+^ cells in glomeruli from *LacZ*/saline, *LacZ*/ADR and *Tmsb4x* treated mice. White arrowheads indicate podocyte nuclei. White dashed line indicates glomerular tuft boundary. Quantification of **(b)** number of WT1^+^ cells in glomeruli and **(c)** glomerular WT1^+^ density. 50 glomeruli were analysed per sample. Quantification of **(d)** WT1^+^ cell count and **(e)** glomerular WT1^+^ density when analysing individual glomeruli. At least 250 glomeruli were analysed per condition. **(f)** Representative images of glomeruli from *LacZ*/saline, *LacZ*/ADR and *Tmsb4x* treated mice immunostained to visualise synaptopodin and F-actin. White arrows indicate F-actin in synaptopodin^+^ areas. Images have been edited to crop out positive staining outside of the glomerular tuft in aid of the macro used for analysis. Quantification of **(g)** mean synaptopodin^+^ F-actin fluorescence, **(h)** area of synaptopodin^+^ covered by F-actin (µm^2^) and **(i)** percentage of synaptopodin^+^ area that was F-actin^+^. Scale bars = 20 µm and the white dashed line indicates glomerular tuft boundaries. Each group contained at least 5 mice and 50 glomeruli were analysed per mouse. Data are presented as mean±SEM. *** P ≤ 0.001. TB4, *Tmsb4x*, thymosin β4; ADR, Adriamycin; WT1, Wilms tumour 1; *LacZ*, β-Galactosidase.

Alterations to podocyte F-actin have been associated with foot process effacement and albuminuria *in vivo* ^9,10^. We hypothesised that the protective effect of TB4 in ADR injury may be partly mediated by its ability to sequester G-actin and regulate F-actin polymerisation.^13,14,46^ Using synaptopodin as a podocyte marker,^47^ and phalloidin to visualise F-actin filaments, we quantified F-actin in the synaptopodin-positive regions (**Figure 4f**). We found that there was no difference in the mean podocyte F-actin fluorescence between any of the groups (**Figure 4g**). There were also no changes to the total area (μm^2^) or percentage of podocyte area covered by F-actin, which remained at approximately 65% (**Figure 4h, i**), indicating that neither ADR nor TB4 alter the amount of F-actin within podocytes.

Albumin leakage can also occur via transcellular transport mediated by endocytosis of albumin in podocyte vesicles followed by excretion to the urinary space.^48–51^ Double staining of glomeruli with synaptopodin and F-actin revealed vesicle-like structures with a diameter of 1-1.3μm within podocytes **(Figure 5a-c)**. There was a significant increase in the number of podocyte vesicles per glomerulus in *LacZ*/ADR (7.17 ±0.32) compared with *LacZ*/saline treated mice (5.33 ±0.26; P<0.001). TB4 treatment prevented this increase (5.58 ±0.26 vesicles per glomerulus; P<0.01) **(Figure 5d)**. To account for changes in glomerular area, vesicle number was normalised to glomerular area (vesicle density). *LacZ*/saline glomeruli had a vesicle density of 2.29 ⨯10-3 ±0.11 ⨯10^−3^ vesicles per μm^2^ which increased to 2.71 ⨯10^−3^ ±0.11 vesicles per μm^2^ after ADR treatment (P<0.001). *Tmsb4x*/ADR treated glomeruli had a significantly lower vesicle density than *LacZ*/ADR glomeruli (2.24 ⨯10-3 ±0.09 ⨯10^−3^ vesicles per μm^2^; P<0.05) **(Figure 5e)**.

**Figure 5.**
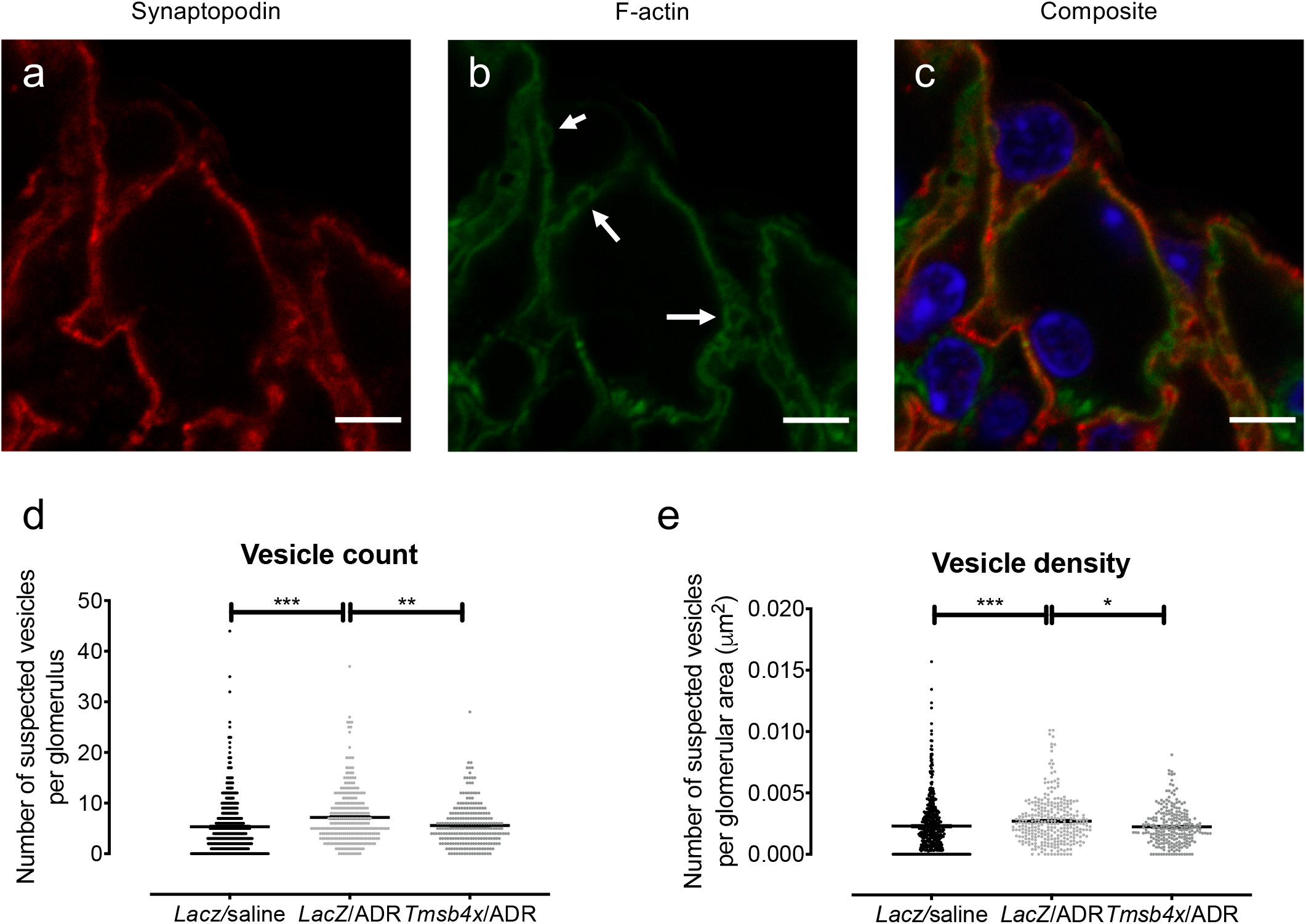
TB4 prevents ADR-induced podocyte vesicle formation. Representative images of **(a)** synaptopodin, **(b)** F-actin and **(c)** the composite including nuclei stained with Hoechst 33342 from a *LacZ*/ADR treated glomerulus. White arrows indicate vesicles. Scale bar = 5 µm. Quantification of **(d)** the number of vesicles per glomerulus and **(e**) vesicle density. Each group contained at least 5 mice and 50 glomeruli were analysed per mouse. Data are presented as mean±SEM. *P≤0.05, **P≤0.01, ***P≤0.001. TB4, *Tmsb4x*, thymosin β4; ADR, Adriamycin; *LacZ*, β-Galactosidase.

### TB4 prevents ADR-induced podocyte F-actin reorganisation *in vitro*

As we have previously shown that lack of endogenous TB4 affects the organisation of F-actin fibres in cultured podocytes^16^, we next sought to perform a detailed assessment of podocyte F-actin architecture *in vitro*. Cultured differentiated mouse podocytes, an established model to study the regulation of the podocyte cytoskeleton^52,53^ were treated with a low (0.0125 μg/ml) or high (0.125 μg/ml) dose of ADR and the potential of synthetic TB4 (100 ng/ml) to abrogate the effects of ADR was assessed (**Figure 6a**). ADR treatment reduced *Tmsb4x* expression in cultured podocytes which was significant at the high dose (60% decrease; P<0.01; **Figure 6b**) mirroring our findings *in vivo*. Treatment with 0.125 μg/ml of ADR led to a significant decrease in podocyte viability (P<0.05) and podocyte cell area (P<0.01) compared with podocytes treated with media alone, which was not prevented by co-administration of exogenous TB4 (**Figure 6c, d**). Next, we quantified podocyte F-actin. Treatment with ADR did not alter podocyte mean F-actin fluorescence, which remained unaffected by TB4 (**Figure 6e**), in agreement with our *in vivo* finding that neither ADR nor TB4 alter the amount of F-actin within podocytes.

**Figure 6.**
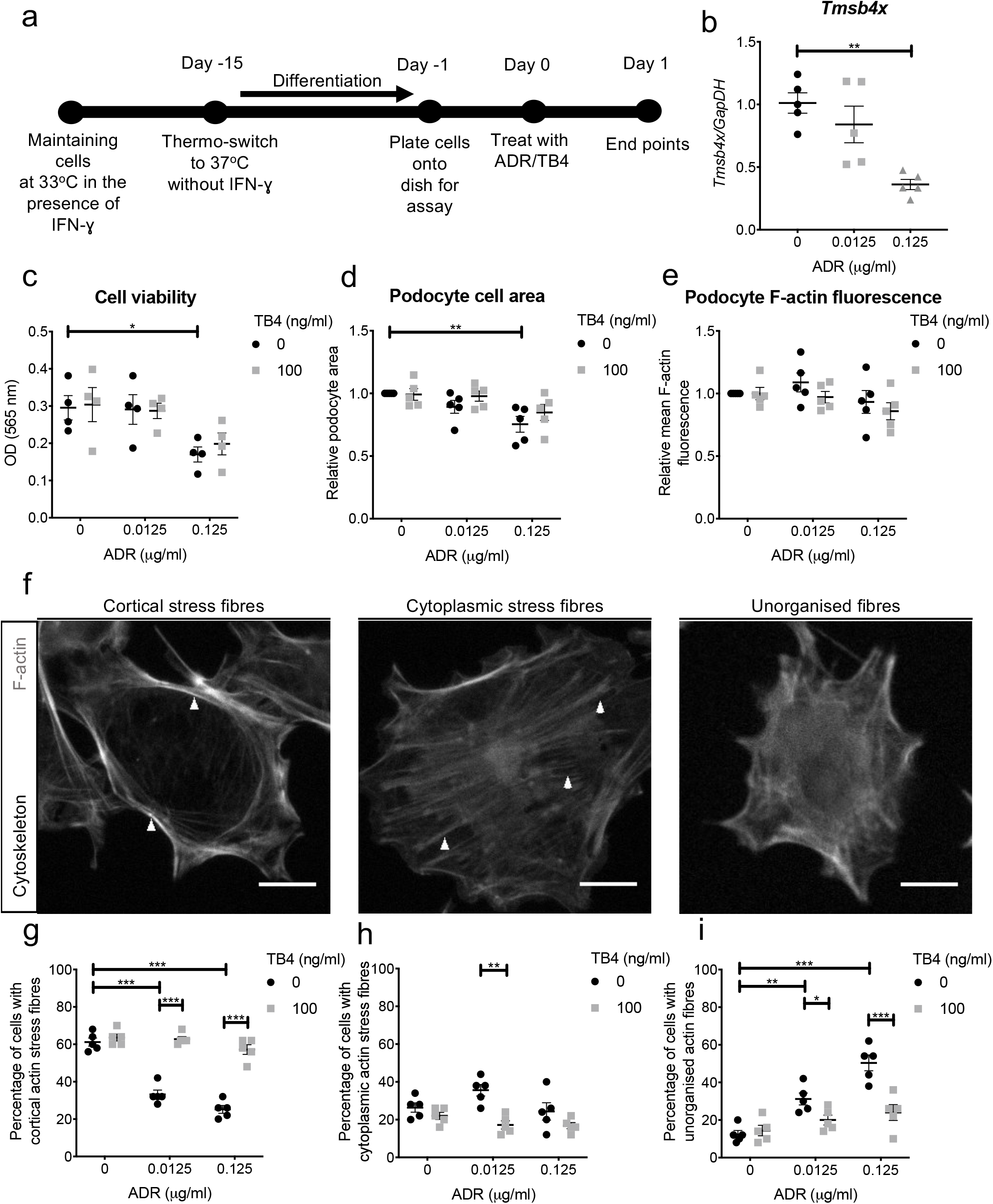
Effect of exogenous TB4 on ADR-injured podocytes *in vitro*. **(a)** Conditionally immortalised mouse podocytes were treated with RPMI-1640/ADR/TB4 and analysed after 24 hours. **(b)** Expression of podocyte *Tmsb4x* mRNA after ADR/TB4 treatment. Effect of ADR/TB4 on **(c)** cell viability, **(d)** podocyte cell area and **(e)** podocyte mean F-actin fluorescence. **(f)** Representative images of podocytes stained with Acti-Stain™ 488 Phalloidin displaying cortical actin stress fibres, cytoplasmic stress fibres and unorganised actin fibres. Scale bar = 10 μm. Percentage of podocytes displaying **(h)** cortical actin stress fibres, **(i)** cytoplasmic stress fibres and **(j)** unorganised stress fibres. All experiments were repeated a minimum of 4 times. For F-actin analysis 50 cells were analysed per experiment \**P* ≤ 0.05, \*\**P* ≤ 0.01 and \*\*\**P* ≤ 0.001. TB4, *Tmsb4x*, thymosin β4; ADR, Adriamycin; IFN-lll, interferon-gamma; OD, optical density; GapDH, Glyceraldehyde 3-phosphate dehydrogenase; MTT, methyltetrazolium.

Finally, to study podocyte F-actin organisation in more detail, we classified F-actin arrangements into cortical actin stress fibres, cytoplasmic stress fibres or unorganised fibres (see examples in **Figure 6f**). Prior to ADR administration, the majority of podocytes (61.2 ±2.2%) had a prevalence of cortical actin stress fibres, compared with 26.4 ±2.5% podocytes with cytoplasmic stress fibres and 12.4 ±2.0% podocytes with unorganised actin fibres. Treatment with 0.0125 μg/ml of ADR changed this distribution with significantly reduced cortical stress fibre prevalence to 33.2 ±2.3% of podocytes (P<0.001) and increased prevalence of unorganised actin fibres (31.2 ±3.2% of podocytes, P<0.01) compared with untreated podocytes. Treatment with exogenous TB4 prevented the ADR-induced F-actin reorganisation and significantly increased the proportion of podocytes with cortical actin stress fibres (62.8 ± 1.4%, P<0.001) and decreased the proportion of podocytes with unorganised actin fibres (20 ±2.6%, P<0.05) compared with the group treated with low dose ADR. Treatment with 0.125 μg/ml of ADR led to exacerbated cytoskeletal disorganisation, reducing cortical stress fibre prevalence to 25.2 ±2.1% of podocytes (P<0.001) and increasing unorganised actin fibre prevalence to 50.4 ±4.2% of podocytes (P<0.001) compared with the untreated group. Co-treatment with exogenous TB4 ameliorated the effects of ADR with cortical stress fibre frequency at 57.2 ±2.6% (P<0.001) and unorganised actin fibre frequency at 24 ±4.2% of podocytes (P<0.001) (**Figure 6g-i**).

## Discussion

In this study we have shown that ADR injury results in reduced *Tmsb4x* mRNA levels in glomeruli and particularly in podocytes. Systemic upregulation of TB4 using AAV-mediated gene therapy prevents ADR-induced albuminuria and podocyte loss *in vivo* and treatment with synthetic TB4 prevents ADR-induced cytoskeletal disorganisation *in vitro*. Thus, we have provided the first evidence that exogenous TB4 can protect podocytes from injury and improve glomerular disease.

Previous studies have demonstrated the expression of endogenous *Tmsb4x* in mouse glomeruli predominately in podocytes.^16,54,55^ The effect of glomerular disease on TB4 levels, however, is less clear. A proteomic study using the rat kidney remnant model of renal fibrosis found that TB4 levels increased threefold in sclerotic versus normal glomeruli.^56^ Our group previously demonstrated that *Tmsb4x* levels were not altered in whole kidneys obtained from mice with glomerulonephritis or in glomerular extracts obtained from human biopsy specimens from patients with rapidly progressive glomerulonephritis or lupus nephritis.^16^ These studies, however, did not assess *Tmsb4x* levels in a cell type-specific manner. Here, we have shown that ADR injury results in a transient reduction of *Tmsb4x* levels in glomeruli and we have performed analysis of a scRNAseq dataset^37^ to demonstrate reduced *Tmsb4x* levels specifically in podocytes.

Since endogenous TB4 has a protective role in glomerular disease,^16,42^ we hypothesised that treatment with exogenous TB4 would be beneficial in ADR injury. Indeed, we found that TB4 administration prevented the onset of albuminuria in mice injured with ADR. Podocyte cells are a crucial component of the glomerular filtration barrier and they are the primary target of ADR injury in the kidney.^33^ We demonstrated that TB4 prevented podocyte loss in ADR-injured mice. Loss of podocytes from the glomerular tuft may result from cell death or from injury that causes podocyte detachment.^2^ The actin cytoskeleton is critical to maintain podocyte shape^57^ and attachment to the GBM.^8^ We developed a novel method to quantify podocyte F-actin *in vivo* and found that neither ADR nor TB4 affected the amount of F-actin in the podocytes. However, detailed analysis of the F-actin cytoskeleton in cultured podocytes revealed that even though there were no changes in the amount of F-actin, ADR injury increased the proportion of podocytes with unorganised actin and this was prevented by treatment with TB4. These findings demonstrate that TB4 protects the podocyte cytoskeleton, prevents podocyte injury and loss which is associated with an improvement in albuminuria following ADR injury. In the epidermis, lack of endogenous TB4 results in hindered eyelid closure and hair follicle angling and defects in planar cell polarity (PCP) with impaired stability of adherens junctions, aberrant F-actin distribution and changes in cell shape.^58^ PCP is also implicated in podocyte health in development and disease.^59^ Van Gogh-like 2 (Vangl2), a core PCP protein, is required for the normal differentiation of glomeruli^60,61^ and podocyte-specific deletion of Vangl2 exacerbates experimental glomerulonephritis in mice.^62,63^ It is therefore possible that some of the effects of TB4 on podocyte cells might be mediated via PCP pathways.

Another mechanism that contributes to albuminuria is the endocytosis of albumin in podocyte vesicles and subsequent excretion to the urinary space. This mechanism has been demonstrated by confocal, electron and intravital multiphoton microscopy in rat models of glomerular injury^48–51^ and has been estimated to contribute 10% of total albumin filtration.^49^ We observed podocyte vesicle-like structures with a diameter of 1-1.3μm, consistent with the size of vesicles reported in the aforementioned studies. These vesicles were more prevalent in ADR-injured glomeruli and their occurrence was reduced by TB4 treatment. Local actin polymerisation is critical for the formation of endocytic vesicles.^64^ TB4 gene therapy may reduce the formation of endocytic vesicles by preventing local actin polymerisation. Indeed, it has been demonstrated that excess concentration of TB4 inhibits endocytosis *in vitro*.^65^ A recent study has demonstrated that TB4 binds to the endocytic receptor, low density lipoprotein receptor related protein 1 (LRP1), and regulates whether endocytosed protein complexes are recycled to the cell surface or targeted for lysosomal degradation.^66^ It remains to be investigated whether TB4 may interact with receptors that facilitate albumin endocytosis and downstream processing in the kidney. The regulation of vesicle formation and transcytosis of albumin by exogenous TB4 may thus represent a novel strategy to limit albuminuria in glomerular disease.

Previous studies have shown that endogenous and exogenous TB4 can improve inflammation in animal models of kidney injury including nephrotoxic nephritis,^16^ angiotensin-II induced hypertensive nephropathy^42^ and acute ischaemia reperfusion injury.^25^ In our study, we assessed the effects of TB4 in the early stages of ADR injury when macrophage infiltration is not present, demonstrating that the protective effect of TB4 is likely independent of its anti-inflammatory properties in this case.

In this study, we used AAV-mediated systemic gene therapy to achieve long term transgene upregulation.^38^ Previous studies have used administration of TB4 protein, which maintains enhanced circulating TB4 levels for only 6 hours.^26^ Our strategy circumvents the quick turnover of TB4 thus achieving sustained levels of upregulation five weeks after AAV administration. Future studies could target TB4-encoding AAVs to the kidney, however, systemic administration of AAV serotypes 1-9 has shown no efficient transduction in the kidney.^38^ Transcriptional targeting,^67^ synthetic AAVs^68^ and novel administration routes, such as administration by retrograde ureteral and subcapsular injections^69^ or by injection into the renal vein,^70^ have shown promise and they could be utilised to achieve kidney-specific overexpression of TB4. Inducible AAVs^71,72^ would enable regulation of the timing of TB4 upregulation to assess its ability to improve the progression of established glomerular disease.

In summary, we have shown that ADR injury results in reduced levels of endogenous TB4, podocyte loss and proteinuria. Systemic gene therapy with TB4 prevents proteinuria and podocyte loss, likely by protecting the actin cytoskeleton and limiting albumin transcytosis. These findings suggest that treatment with TB4 could be a potential novel therapeutic strategy to prevent podocyte injury and maintain filtration in glomerular disease.

## Materials and methods

### scRNAseq analysis

scRNAseq analysis was performed using RStudio for Macintosh (RStudio Inc., v1.2.5042) using R (v4.0.2). The complete annotated R code used to perform the analyses and generate plots used in the manuscript has been deposited in GitHub and can be accessed at https://github.com/davidlonglab/Mason_TB4_2021.

#### Data acquisition

The raw scRNAseq dataset used in this analysis was acquired from a study characterising the single-cell transcriptome of murine ADR nephropathy using the 10X Genomics platform^37^. Matrices of gene counts per droplet, generated after alignment of reads to genes, were acquired from the National Center for Biotechnology Information Gene Expression Omnibus (GSE146912) and are available at https://www.ncbi.nlm.nih.gov/geo/query/acc.cgi?acc=GSE146912.

#### Quality control, data processing and integration

All the following analysis was performed using the Seurat toolkit^73^. The count matrices from n = 2 control samples (8,412 cells) and n = 2 samples with Adriamycin nephropathy (8,296 cells) were merged into a single object. Genes expressed in two or fewer droplets were excluded and droplets with < 200 and > 4000 detected genes and > 10% of features mapping to the mitochondrial genome were excluded. The counts were then normalized using the NormalizeData function and scaled by all detected genes using the ScaleData function before principal component analysis (PCA), using the top nine components for downstream analyses. Integration and matching of cell types between experimental conditions was performed using the Harmony package for R^74^.

#### Clustering, cell type identification and counting

Shared nearest neighbor graphing was performed using the FindNeighbors function. Unsupervised clustering was performed with the FindClusters function using the Louvain algorithm and a resolution of 0.4, generating 14 transcriptionally distinct clusters, before dimension reduction using Uniform Manifold Approximation and Projection (UMAP). Cell type identification was performed by assessing the top ten differentially expressed genes per cluster calculated using the FindAllMarkers function and canonical markers for glomerular cell types were compared from previous scRNAseq studies.^37,75,76^ By grouping clusters with a common cell identity together, ten glomerular cell types were subsequently identified and assigned. The number of cell types by experimental condition was exported and graphed in Prism (GraphPad, v9.0.0)

#### Comparison of Tmsb4x expression

The FindAllMarkers function was used to compare the scaled expression of *Tmsb4x* between ADR injury and control datasets. The average log fold change was calculated for podocytes or all glomerular cell types (glomerular endothelial cells, mesangial cells and podocytes) between experimental conditions. Wilcoxon Rank Sum tests was used to assess statistical significance, with an adjusted P value of ≤ 0.05.

### Adeno-associated viral generation

The recombinant AAV-*Tmsb4x* and AAV-*LacZ* vectors were produced using triple transfection in HEK293 cells. Cells were harvested and virus was purified by iodixanol-gradient centrifugation. The virus was further purified using Sepharose G100 SF resin (Merck, Darmstadt, Germany) in Econopac colums (Bio-Rad, Watford, UK). Virus was concentrated in PBS using Amicon Ultra-15 Centrifugal Filter Units (Merck) and stored at 4°C.^31,77,78^ Viral titre was quantified by inverted terminal repeat probe quantitative polymerase chain reaction (PCR). Helper plasmid delta F6 was purchased from Puresyn (Malvern, PA).

### Experimental animals and procedures

All experiments were carried out according to a UK Home Office project and were compliant with the UK Animals (Scientific Procedures) Act 1986. Male BALB/c mice aged between 7-10 weeks were administered with AAV (sub-serotype 2/7; 5⨯10^12^ viral particles per mouse) expressing *LacZ* (AAV-*LacZ*) or *Tmsb4x* (AAV-*Tmsb4x*) via the tail vein. To induce glomerular injury, mice were intravenously injected with 10 mg/kg of ADR (Merck) or vehicle (0.9 % saline) 21 days after AAV administration.

### Renal function

Urine was collected from mice by housing them individually in metabolic cages overnight. Blood samples were collected from the lateral saphenous vein. Albumin concentrations were measured by enzyme-linked immunosorbent assay^36,79^ (Bethyl Laboratories, Montgomery, TX). Urinary creatinine concentrations were quantified using a commercially available kit (Cayman Chemicals, Ann Arbor, MI). Blood urea nitrogen was assessed using a commercially available assay kit validated in mice (BioAssay Systems, Hayward, CA).^80^

### TB4 enzyme linked immunosorbent assay (ELISA)

The plasma concentration of TB4 was determined by ELISA based on the protocol previously described by Mora et. al.^26^ Standards of 10,000 ng/ml, 5,000 ng/ml, 2,500 ng/ml, 1,250 ng/ml, 625 ng/ml, 312.5 ng/ml, 156 ng/ml, 78 ng/ml and 39 ng/ml TB4 were prepared using synthetic TB4 (ReGeneRx Biopharmaceuticals Inc, Rockville, MD) diluted in incubation buffer ((pH 7.4, Na_2_HPO_4_ (0.01M), NaCl (0.15M), Tween-20 (0.055 % v/v), BSA (1 % v/v)). Equal volumes of standards or samples, incubation buffer and TB4 antibody prediluted at 1:4000 (AF6796; R&D Systems, Minneapolis, MN,) were added to sterile borosilicate tubes and incubated overnight at 4 °C. A flat bottom 96 well plate was coated with 100 μl of 50 ng/ml recombinant TB4 in carbonate-bicarbonate buffer and incubated overnight at 4°C. Negative control wells were coated with buffer only. The plate was washed with washing buffer ((pH 7.4, Na_2_HPO_4_ (0.01M), NaCl (0.15M), CaCl_2_ (1 mM), MgCl_2_ (0.5 mM), Tween-20 (0.55% v/v)), blocked with 200 μl of blocking buffer (5 % dry fat milk in incubation buffer) and 100 μl of each standard and sample were added to the appropriate wells and incubated for 2 hours. Following washing, 100 μl of goat anti-sheep HRP-conjugated secondary antibody (61-8620, Thermo Fisher Scientific) diluted 1:2000 in incubation buffer was added to each well and incubated for 1 hour before washing. Substrate solution (100 μl per well) containing equal parts stabilised H_2_O_2_ and stabilised tetramethylbenzidine was added and 15 minutes later the reaction was stopped with the addition of 50 μl of 2M sulphuric acid per well and absorbance was read at 450 nm using a plate reader (M200 Pro, Tecan, Männedorf Switzerland).

### Tissue processing and immunostaining

Tissues were fixed in 4% paraformaldehyde in PBS. To prepare wax sections, tissues were dehydrated, embedded in paraffin and 5 μm thick sections were prepared. To prepare cryosections, tissues were placed overnight in 30% sucrose in PBS, embedded in Tissue-Tek optimal cutting temperature compound (Agar Scientific, Stansted, UK) and 8 μm sections were prepared.

Immunohistochemistry was performed for TB4 (AF6796, R&D Systems) followed by secondary rabbit anti-sheep antibodies (Thermo Fisher Scientific, Waltham, MA) and ImmPRESS polymer anti-rabbit IgG reagent (Vector Laboratories, Burlingame, CA) conjugated to horseradish peroxidase and detected by 3,3’-diaminobenzidine. Images were obtained on a Leica DM5500 B brightfield microscope (Leica Biosystems, Milton Keynes, UK).

Immunofluorescence was performed^81^ using primary antibodies against WT1 (AB89901, Abcam, Cambridge, UK), synaptopodin (163-004-SY, Synaptic Systems, Goettingen, Germany) and F4/80 (MCA497R, Bio-Rad), followed by appropriate AlexaFluor594 and AlexaFluor488 (Thermo Fisher Scientific, Waltham, MA) secondary antibodies. Negative controls consisted of omission of primary antibodies. Acti-Stain 488™ Phalloidin (Cytoskeleton, Denver, CO) was used to visualise actin filaments. Images were acquired using a Zeiss Laser Scanning 880 confocal microscope with a 63x NA1.4 Oil Plan Apochromat objective (Carl Zeiss, Oberkochen, Germany).

The number of WT1^+^ cells found within the glomerular tuft was counted in 50 glomeruli per mouse. Podocyte density was calculated as the number of WT1^+^ cells in the glomerular tuft, normalized to the glomerular area (measured using ImageJ^82^). F4/80^+^ cells within the glomerular tuft and in the peri-glomerular area were counted in 50 glomeruli per mouse. Podocyte vesicles were identified as circular structures that were surrounded by F-actin and were between 1 and 1.3 μm in diameter. The number of vesicles and vesicle density (number of vesicles/glomerular area) was counted in 50 glomeruli per mouse. For automated quantification of podocyte F-actin, a macro was generated, which has been deposited in GitHub (https://github.com/DaleMoulding/Fiji-Macros/blob/master/README.md#podocyte-f-actin--synaptopodin). A Gaussian blur with a Sigma (radius) value of 2.0 was applied to each channel to create a solid mask of synaptopodin to outline the podocyte area. The mean fluorescence of F-actin within the synaptopodin positive area was measured along with the total area (μm^2^) and percentage area of F-actin in the synaptopodin positive regions.

### Cell culture

Mouse podocytes^52^ were cultured as described^60^ and allowed to differentiate for 14 days. Cells were treated with 100 ng/ml synthetic TB4 (ReGeneRx Biopharmaceuticals Inc) and either a low (0.0125 μg/ml) or a high (0.125 μg/ml) dose of ADR for 24 hours.

Cell viability was determined by the methyltetrazolium assay. To visualize F-actin filaments, podocytes were fixed in 4% paraformaldehyde and 4% sucrose and stained with Acti-stain™ 488 Phalloidin (Cytoskeleton) and 50 cells per condition were assessed. The area of each cell and the mean F-actin fluorescence were quantified using ImageJ. Actin filaments were classified as cortical stress fibers, cytoplasmic stress fibers or unorganised actin.

### Quantitative real-time PCR

RNA extracted from mouse whole-kidney (500 ng), glomerular extracts (100 ng; isolated by Dynabeads^36^ (Thermo Fisher Scientific) or cultured podocytes (100 ng) was used to prepare cDNA (iScript kit, Bio-Rad), and quantitative real-time PCR was performed as described previously^36^ with GapDH as a housekeeping gene. All measurements were performed in duplicate.

### Statistical analysis

All samples were assessed by independent observers blinded to treatment group. Data are presented as mean ± SEM and were analysed using GraphPad Prism v9 (GraphPad Software, La Jolla, CA). Normal distribution was assessed by Shapiro-Wilk test. When differences between 2 groups were evaluated, data were analysed using a t test. When 3 or more groups were assessed, 1-way ANOVA with Tukey’s multiple comparison post hoc tests were used. Data affected by 2 variables were analysed using 2-way ANOVA with Tukey’s multiple comparison post hoc tests. For analysis of individual glomeruli, data were analysed by Kruskal-Wallis non-parametric test followed by Dunn’s multiple comparisons. Statistical significance was accepted at P ≤ 0.05.

## Acknowledgements

All mice were maintained by staff at GOSICH Western Laboratories and UCL Biological Services. Microscopy was performed at the University of Greenwich Imaging Facility. Synthetic TB4 was provided by ReGeneRx Biopharmaceuticals Inc, Rockville, MD. This work was supported by a Kidney Research UK Intermediate Fellowship (PDF8/2015 to EV), a University of Kent VC Ph.D. Studentship (to EV), a Wellcome Trust Postdoctoral Training Fellowship for MB/Ph.D. graduates (095949/Z/11/Z, to EP), project grants from the Medical Research Council (MR/P018629/1 and MR/J003638/1, to DAL), a Child Health Research Ph.D. Studentship from UCL Great Ormond Street Institute of Child Health (to DJ, DAL) and a Diabetes UK studentship (17/0005733 to DAL). Professor Long’s laboratory is supported by the NIHR Biomedical Research Centre at Great Ormond Street Hospital for Children NHS Foundation Trust and University College London.

## Author contributions

Conceptualisation, EV, WJM and DAL; Methodology, WJM, EV, EP, SP, AW; Software and Data Curation, DJJ, GP and DAM; Formal Analysis, WJM, DJJ, GP and EV; Investigation, WJM, EV, MK-J, EP and SP; Writing -Original Draft, WJM, DAL and EV; Writing -Review & Editing, WJM, DJJ, CK, CP-W, EP, PRR, DAL and EV; Resources, CK; Supervision, EV, DAL and CP-W; Funding Acquisition EV and DAL

## References

1. Bikbov B, Purcell CA, Levey AS et al. Global, regional, and national burden of chronic kidney disease, 1990–2017: a systematic analysis for the Global Burden of Disease Study 2017. Lancet 2020; 395: 709–733.

2. Greka A, Mundel P. Cell biology and pathology of podocytes. Annu. Rev. Physiol. 2012; 74: 299–323.

3. Miner JH. Glomerular basement membrane composition and the filtration barrier. Pediatr. Nephrol. 2011; 26: 1413–1417.

4. Pavenstädt H, Kriz W, Kretzler M. Cell Biology of the Glomerular Podocyte. Physiol. Rev. 2003; 83: 253–307.

5. Reiser J, Altintas MM. Podocytes. F1000Research 2016; 5:F1000.

6. Welsh GI, Saleem MA. The podocyte cytoskeleton-Key to a functioning glomerulus in health and disease. Nat. Rev. Nephrol 2012; 8: 14–21.

7. Ichimura K, Kurihara H, Sakai T. Actin Filament Organization of Foot Processes in Rat Podocytes. J. Histochem. Cytochem. 2003; 51: 1589–1600.

8. Sachs N, Sonnenberg A. Cell-matrix adhesion of podocytes in physiology and disease. Nat. Rev. Nephrol 2013; 9: 200–210.

9. Suleiman HY, Roth R, Jain S et al. Injury-induced actin cytoskeleton reorganization in podocytes revealed by super-resolution microscopy. JCI insight 2017; 2(16):e94137

10. Yu H, Suleiman H, Kim AHJ et al. Rac1 activation in podocytes induces rapid foot process effacement and proteinuria. Mol. Cell. Biol. 2013; 33: 4755–4764.

11. Harvey SJ, Jarad G, Cunningham J et al. Podocyte-Specific Deletion of Dicer Alters Cytoskeletal Dynamics and Causes Glomerular Disease. J Am Soc Nephrol 2008; 19: 2150– 2158.

12. Benzing T, Salant D. Insights into Glomerular Filtration and Albuminuria. Ingelfinger JR, ed. N. Engl. J. Med. 2021; 384: 1437–1446.

13. Sanders MC, Goldstein AL, Wang Y-L. Thymosin B4 (Fx peptide) is a potent regulator of actin polymerization in living cells. Cell Biol. 1992; 89: 4678–4682.

14. Safer D, Elzinga M, Nachmias VT. Thymosin B4 and Fx, an Actin-sequestering Peptide, Are Indistinguishable*. J. Biol. Chem. 1991; 266: 4029–4032.

15. Xue B, Leyrat C, Grimes JM et al. Structural basis of thymosin-β4/profilin exchange leading to actin filament polymerization. Proc. Natl. Acad. Sci. 2014; 111: E4596–E4605.

16. Vasilopoulou E, Kolatsi-Joannou M, Lindenmeyer MT et al. Loss of endogenous thymosin β4 accelerates glomerular disease. Kidney Int. 2016; 90: 1056–1070.

17. Smart N, Bollini S, Dubé KN et al. De novo cardiomyocytes from within the activated adult heart after injury. Nature 2011; 474: 640–644.

18. Sosne G, Szliter EA, Barrett R et al. Thymosin Beta 4 Promotes Corneal Wound Healing and Decreases Inflammation in Vivo Following Alkali Injury. Exp. Eye Res. 2002; 74: 293–299.

19. Morris DC, Cui Y, Cheung WL et al. A dose–response study of thymosin β4 for the treatment of acute stroke. J. Neurol. Sci. 2014; 345: 61–67.

20. Conte E, Genovese T, Gili E et al. Thymosin β4 protects C57BL/6 mice from bleomycin-induced damage in the lung. Eur. J. Clin. Invest. 2013; 43: 309–315.

21. Vasilopoulou E, Riley PR, Long DA. Thymosin-β4: A key modifier of renal disease. Expert Opin. Biol. Ther. 2018; 18: 185–192.

22. Zhu J, Su L-P, Zhou Y et al. Thymosin β4 Attenuates Early Diabetic Nephropathy in a Mouse Model of Type 2 Diabetes Mellitus. Am. J. Ther. 2015; 22: 141–146.

23. Yuan J, Shen Y, Yang X et al. Thymosin β4 alleviates renal fibrosis and tubular cell apoptosis through TGF-β pathway inhibition in UUO rat models. BMC Nephrol. 2017; 18: 314.

24. Zuo Y, Chun B, Potthoff SA et al. Thymosin beta4 and its degradation product, Ac-SDKP, are novel reparative factors in renal fibrosis. Kidney Int 2013; 84: 1166–1175.

25. Aksu U, Yaman OM, Guner I et al. The Protective Effects of Thymosin-β-4 in a Rat Model of Ischemic Acute Kidney Injury. J. Investig. Surg. 2019; 8: 1–9.

26. Mora CA, Baumann CA, Paino JE et al. Biodistribution of synthetic thymosin beta 4 in the serum, urine and major organs of mice. Int. J. Immunopharmac 1997; 19: 1–8.

27. Weber M, Rabinowitz J, Provost N et al. Recombinant adeno-associated virus serotype 4 mediates unique and exclusive long-term transduction of retinal pigmented epithelium in rat, dog, and nonhuman primate after subretinal delivery. Mol. Ther. 2003; 7: 774–781.

28. Nathwani AC, Tuddenham EGD, Rangarajan S et al. Adenovirus-Associated Virus Vector– Mediated Gene Transfer in Hemophilia B. N. Engl. J. Med. 2011; 365: 2357–2365.

29. Wang D, Tai P, Gao G. Adeno-associated virus vector as a platform for gene therapy delivery. Nat. Rev. Drug Discov. 2019; 18: 358–378.

30. Ziegler T, Bähr A, Howe A et al. Tβ4 Increases Neovascularization and Cardiac Function in Chronic Myocardial Ischemia of Normo-and Hypercholesterolemic Pigs. Mol. Ther. 2018; 26: 1706–1714.

31. Hinkel R, Trenkwalder T, Petersen B et al. MRTF-A controls vessel growth and maturation by increasing the expression of CCN1 and CCN2. Nat Commun 2014; 5: 3970.

32. Bongiovanni D, Ziegler T, D’Almeida S et al. Thymosin beta4 attenuates microcirculatory and hemodynamic destabilization in sepsis. Expert Opin Biol Ther 2015; 15 Suppl 1: S203–10.

33. Papeta N, Zheng Z, Schon EA et al. Prkdc participates in mitochondrial genome maintenance and prevents Adriamycin-induced nephropathy in mice. J. Clin. Invest. 2010; 120: 4055–4064.

34. Ni Y, Wang X, Yin X et al. Plectin protects podocytes from adriamycin-induced apoptosis and F-actin cytoskeletal disruption through the integrin α6β4/FAK/p38 MAPK pathway. J. Cell. Mol. Med. 2018; 22: 5450–5467.

35. Dai R, Liu H, Han X et al. Angiopoietin-like-3 knockout protects against glomerulosclerosis in murine adriamycin-induced nephropathy by attenuating podocyte loss. BMC Nephrol. 2019; 20: 1–11.

36. Long DA, Kolatsi-Joannou M, Price KL et al. Albuminuria is associated with too few glomeruli and too much testosterone. Kidney Int. 2013; 83: 1118–1129.

37. Chung J-J, Goldstein L, Chen Y-JJ et al. Single-Cell Transcriptome Profiling of the Kidney Glomerulus Identifies Key Cell Types and Reactions to Injury. J. Am. Soc. Nephrol. 2020; 31: 2341–2354.

38. Zincarelli C, Soltys S, Rengo G et al. Analysis of AAV serotypes 1-9 mediated gene expression and tropism in mice after systemic injection. Mol Ther 2008; 16: 1073–1080.

39. Wise T, MacDonald GJ, Klindt J et al. Characterization of thymic weight and thymic peptide thymosin-beta 4: effects of hypophysectomy, sex, and neonatal sexual differentiation. Thymus 1992; 19: 235–244.

40. Brinkkoetter PT, Ising C, Benzing T. The role of the podocyte in albumin filtration. Nat. Rev. Nephrol. 2013; 9: 328–336.

41. Kirtane AJ, Leder DM, Waikar SS et al. Serum Blood Urea Nitrogen as an Independent Marker of Subsequent Mortality Among Patients With Acute Coronary Syndromes and Normal to Mildly Reduced Glomerular Filtration Rates. J. Am. Coll. Cardiol. 2005; 45: 1781– 1786.

42. Kumar N, Liao T-D, Romero CA et al. Thymosin β4 Deficiency Exacerbates Renal and Cardiac Injury in Angiotensin-II-Induced Hypertension. Hypertens. (Dallas, Tex. 1979) 2018; 71: 1133–1142.

43. Zhong F, Wang W, Lee K et al. Role of C/EBP-α in Adriamycin-induced podocyte injury. Sci. Rep. 2016; 6:33520.

44. Guo J-K, Menke AL, Gubler M-C et al. WT1 is a key regulator of podocyte function: reduced expression levels cause crescentic glomerulonephritis and mesangial sclerosis. Hum. Mol. Genet. 2002; 11: 651–9.

45. Jefferson JA, Shankland SJ. The Pathogenesis of Focal Segmental Glomerulosclerosis. Adv. Chronic Kidney Dis. 2014; 21: 408–416.

46. Hannappel E, Wartenberg F. Actin-sequestering ability of thymosin beta 4, thymosin beta 4 fragments, and thymosin beta 4-like peptides as assessed by the DNase I inhibition assay. Biol. Chem. Hoppe. Seyler. 1993; 374: 117–122.

47. Wang J, Hidaka T, Sasaki Y et al. Neurofilament heavy polypeptide protects against reduction in synaptopodin expression and prevents podocyte detachment. Sci. Rep. 2018; 8: 1–14.

48. Kinugasa S, Tojo A, Sakai T et al. Selective albuminuria via podocyte albumin transport in puromycin nephrotic rats is attenuated by an inhibitor of NADPH oxidase. Kidney Int. 2011; 80: 1328–38.

49. Maria Schießl I, Hammer A, Kattler V et al. Intravital Imaging Reveals Angiotensin II-Induced Transcytosis of Albumin by Podocytes. J Am Soc Nephrol 2016; 27: 731–744.

50. Burford JL, Gyarmati G, Shirato I et al. Combined use of electron microscopy and intravital imaging captures morphological and functional features of podocyte detachment. Pflügers Arch. - Eur. J. Physiol. 2017; 469: 965–974.

51. Ichimura K, Miyaki T, Kawasaki Y et al. Morphological Processes of Foot Process Effacement in Puromycin Aminonucleoside Nephrosis Revealed by FIB/SEM Tomography. J Am Soc Nephrol 2019 30(1): 96–108.

52. Mundel P, Reiser J, Zúñiga Mejía Borja A et al. Rearrangements of the cytoskeleton and cell contacts induce process formation during differentiation of conditionally immortalized mouse podocyte cell lines. Exp. Cell Res. 1997; 236: 248–58.

53. Shankland SJ, Pippin JW, Reiser J et al. Podocytes in culture: past, present, and future. Kidney Int 2007; 72: 26–36.

54. Guinobert I, Viltard M, Piquemal D et al. Identification of differentially expressed genes between fetal and adult mouse kidney: candidate gene in kidney development. Nephron Physiol 2006; 102: 81–91.

55. Brunskill EW, Georgas K, Rumballe B et al. Defining the molecular character of the developing and adult kidney podocyte. PLoS One 2011; 6: e24640.

56. Xu BJ, Shyr Y, Liang X et al. Proteomic patterns and prediction of glomerulosclerosis and its mechanisms. J Am Soc Nephrol 2005; 16: 2967–2975.

57. Sever S, Schiffer M. Actin dynamics at focal adhesions: a common endpoint and putative therapeutic target for proteinuric kidney diseases. Kidney Int. 2018; 93: 1298–1307.

58. Padmanabhan K, Grobe H, Cohen J et al. Thymosin β4 is essential for adherens junction stability and epidermal planar cell polarity. Development 2020; 147: 1–15.

59. Papakrivopoulou E, Jafree DJ, Dean CH et al. The Biological Significance and Implications of Planar Cell Polarity for Nephrology. Front. Physiol. 2021; 12: 599529.

60. Yates LL, Papakrivopoulou J, Long DA et al. The planar cell polarity gene Vangl2 is required for mammalian kidney-branching morphogenesis and glomerular maturation. Hum Mol Genet 2010; 19: 4663–4676.

61. Babayeva S, Rocque B, Aoudjit L et al. Planar cell polarity pathway regulates nephrin endocytosis in developing podocytes. J. Biol. Chem. 2013; 288: 24035–48.

62. Rocque BL, Babayeva S, Li J et al. Deficiency of the planar cell polarity protein Vangl2 in podocytes affects glomerular morphogenesis and increases susceptibility to injury. J. Am. Soc. Nephrol. 2015; 26: 576–86.

63. Papakrivopoulou E, Vasilopoulou E, Lindenmeyer MT et al. Vangl2, a planar cell polarity molecule, is implicated in irreversible and reversible kidney glomerular injury. J. Pathol. 2018; 246: 485–496.

64. Hinze C, Boucrot E. Local actin polymerization during endocytic carrier formation. Biochem. Soc. Trans. 2018; 46: 565–576.

65. Lamaze C, Fujimoto LM, Yin HL et al. The actin cytoskeleton is required for receptor-mediated endocytosis in mammalian cells. J. Biol. Chem. 1997; 272: 20332–5.

66. Munshaw S, Bruche S, Redpath AN et al. Thymosin β4 protects against aortic aneurysm via endocytic regulation of growth factor signaling. J. Clin. Invest. 2021.

67. Schievenbusch S, Strack I, Scheffler M et al. Combined paracrine and endocrine AAV9 mediated expression of hepatocyte growth factor for the treatment of renal fibrosis. Mol. Ther. 2010; 18: 1302–1309.

68. Ikeda Y, Sun Z, Ru X et al. Efficient gene transfer to kidney mesenchymal cells using a synthetic adeno-associated viral vector. J. Am. Soc. Nephrol. 2018; 29: 2287–2297.

69. Rubin JD, Nguyen T V., Allen KL et al. Comparison of Gene Delivery to the Kidney by Adenovirus, Adeno-Associated Virus, and Lentiviral Vectors after Intravenous and Direct Kidney Injections. Hum. Gene Ther. 2019; 30: 1559–1571.

70. Zhong F, Chen H, Xie Y et al. Protein S protects against podocyte injury in diabetic nephropathy. J. Am. Soc. Nephrol. 2018; 29: 1397–1410.

71. Vanrell L, Di Scala M, Blanco L et al. Development of a liver-specific tet-on inducible system for AAV vectors and its application in the treatment of liver cancer. Mol. Ther. 2011; 19: 1245–1253.

72. Chtarto A, Humbert-Claude M, Bockstael O et al. A regulatable AAV vector mediating GDNF biological effects at clinically-approved sub-antimicrobial doxycycline doses. Mol. Ther. - Methods Clin. Dev. 2016; 3: 16027.

73. Butler A, Hoffman P, Smibert P et al. Integrating single-cell transcriptomic data across different conditions, technologies, and species. Nat. Biotechnol. 2018; 36: 411–420.

74. Korsunsky I, Millard N, Fan J et al. Fast, sensitive and accurate integration of single-cell data with Harmony. Nat. Methods 2019; 16: 1289–1296.

75. Karaiskos N, Rahmatollahi M, Boltengagen A et al. A single-cell transcriptome atlas of the mouse glomerulus. J. Am. Soc. Nephrol. 2018; 29: 2060–2068.

76. Fu J, Akat KM, Sun Z et al. Single-cell RNA profiling of glomerular cells shows dynamic changes in experimental diabetic kidney disease. J. Am. Soc. Nephrol. 2019; 30: 533–545.

77. Grieger JC, Choi VW, Samulski RJ. Production and characterization of adeno-associated viral vectors. Nat. Protoc. 2006; 1: 1412–28.

78. Moretti A, Fonteyne L, Giesert F et al. Somatic gene editing ameliorates skeletal and cardiac muscle failure in pig and human models of Duchenne muscular dystrophy. Nat. Med. 2020; 26: 207–214.

79. Dessapt-Baradez C, Woolf AS, White KE et al. Targeted glomerular angiopoietin-1 therapy for early diabetic kidney disease. J Am Soc Nephrol 2014; 25: 33–42.

80. Kolatsi-Joannou M, Price KL, Winyard PJ et al. Modified citrus pectin reduces galectin-3 expression and disease severity in experimental acute kidney injury. PLoS One 2011; 6: e18683.

81. Huang JL, Woolf AS, Kolatsi-Joannou M et al. Vascular Endothelial Growth Factor C for Polycystic Kidney Diseases. J Am Soc Nephrol 2016; 27: 69–77.

82. Schindelin J, Arganda-Carreras I, Frise E et al. Fiji: an open-source platform for biological-image analysis. Nat. Methods 2012; 9: 676–682.

