## Supplementary Data for "Systemic gene therapy with thymosin β4 alleviates glomerular injury in mice"

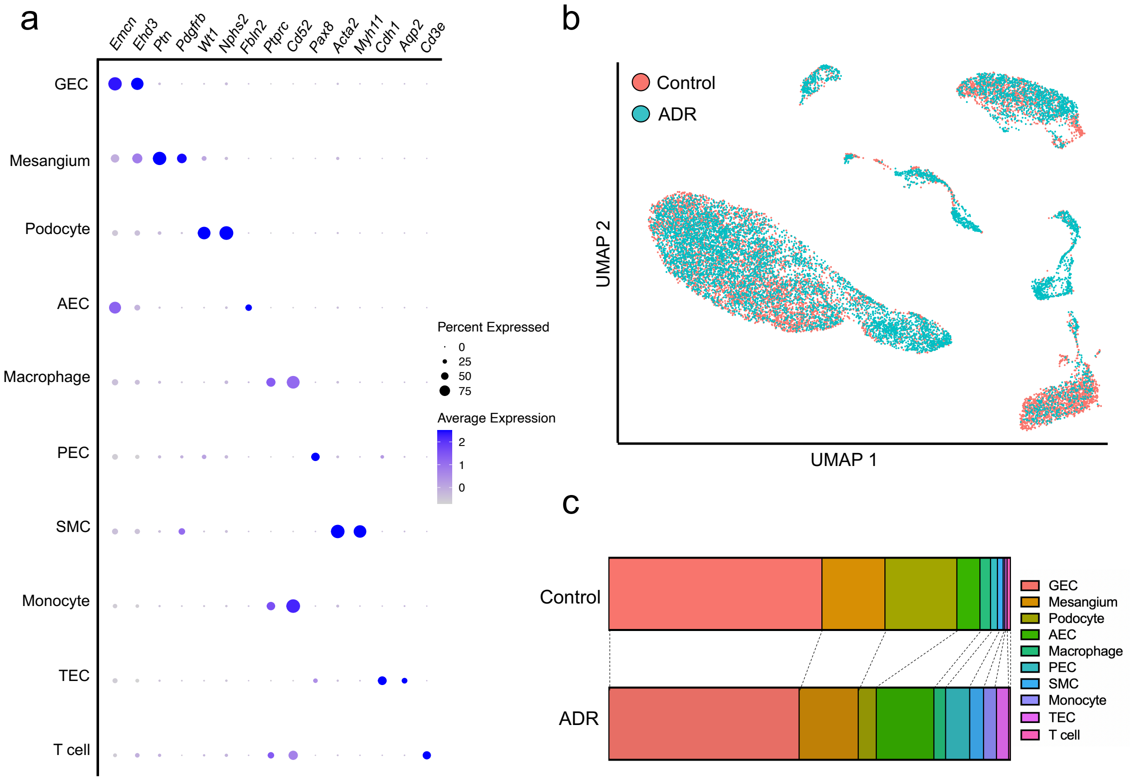


**Supplementary Figure 1. Cell type identification and comparison of cell type proportions within single-cell RNA sequencing data. (a)** Dotplot showing enrichment of canonical markers between cell types within the single-cell RNA sequencing (scRNAseq) dataset. The markers include endomucin (*Emcn)* and Eps15 homology domain-containing protein 3 *(Edh3)* for glomerular endothelial cells (GEC), pleiotrophin *(Ptn)* and platelet-derived growth factor receptor beta *(Pdgfrb)* for mesangial cells, Wilms' tumour 1 *(Wt1)* and podocin (*Nphs2)* for podocytes, *Emcn* and fibulin 2 *(Fbln2)* for arterial endothelial cells (AEC), protein tyrosine phosphatase receptor type C *(Ptprc)* and campath 1 antigen *(Cd52)* for monocytes and macrophages, paired box gene 8 *(Pax8)* for parietal epithelial cells (PEC), actin alpha 2 *(Acta2)* and myosin heavy chain 11 *(Myh11)* for smooth muscle cells (SMC), E-cadherin

*(Cdh1)* and aquaporin 2 *(Aqp2)* for tubular epithelial cells (TEC) and T cell surface glycoprotein CD3 epsilon chain *(Cd3e)* for T cells. **(b)** Uniform manifold approximation and projection (UMAP) grouped by experimental condition in the scRNAseq dataset. The UMAP corresponds to **Figure 1A**, showing concordance of cell types between Adriamycin nephropathy (ADR) and control. **(c)** Bar graphs comparing the proportions of cell types between ADR and control. In the control dataset, *n* = 4,402 GECs, *n* = 1,302 mesangial cells, *n* = 1,486 podocytes, *n* = 477 AECs, *n* = 218 macrophages, *n* = 143 PECs, *n* = 118 SMCs, *n* = 34 monocytes, *n* = 47 TECs and *n* = 69 T cells were detected. In the ADR dataset *n* = 3,895 GECs, *n* = 1,239 mesangial cells, *n* = 378 podocytes, *n* = 1,207 AECs, *n* = 245 macrophages, *n* = 505 PECs, *n* = 290 SMCs, *n* = 271 monocytes, *n* = 256 TECs and *n* = 36 T cells were detected.

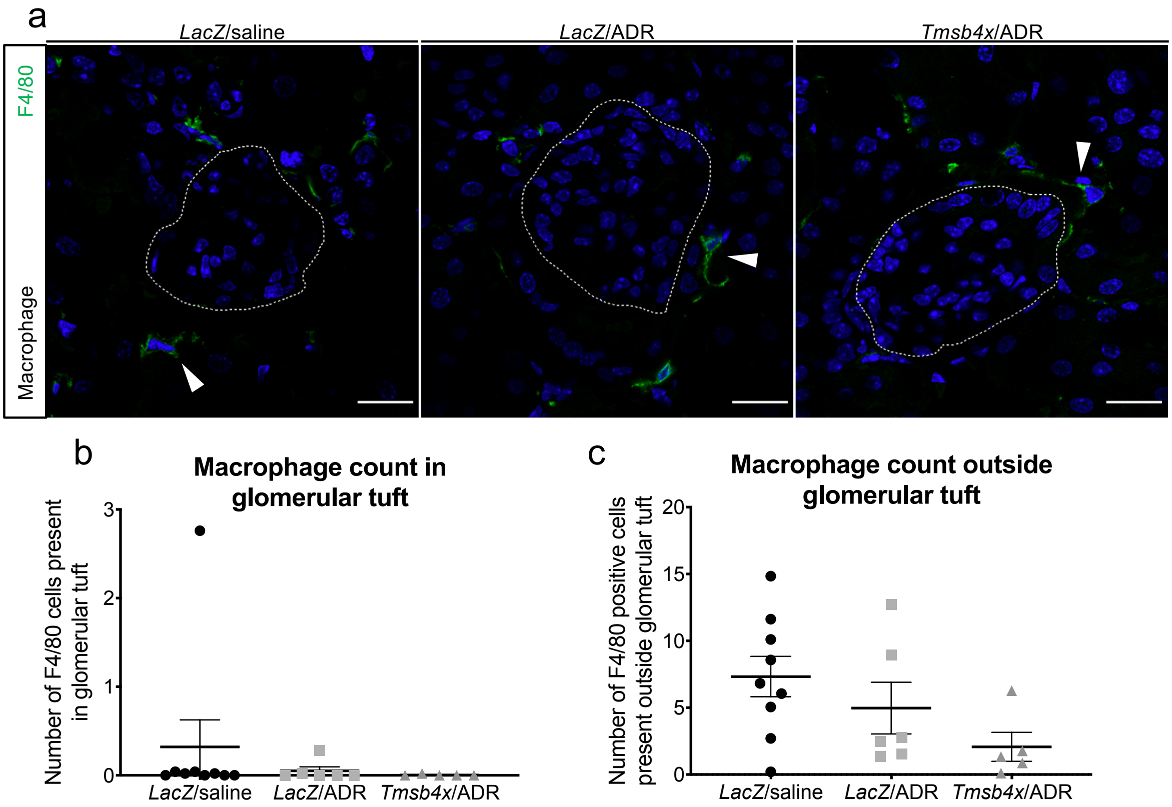


**Supplementary Figure 2. Adriamycin does not cause macrophage infiltration at 14 days.** **(a)** Representative images of F4/80 expression around *LacZ*/saline, *LacZ/*ADR and *Tmsb4x*/ADR glomeruli. White arrowheads indicate positive F4/80 staining. White dashed line indicates the glomerular tuft. Quantification of F4/80 positive cells **(b)** inside the glomerular tuft and **(c)** outside the glomerular tuft. Each group contained a minimum of 5 mice and 50 glomeruli were analysed per mouse. Data are presented as mean±SEM. Scale bars = 20 µm. TB4, *Tmsb4x,* thymosin β4; ADR, Adriamycin; *LacZ*, β-Galactosidase.
